# Modulation of spinal circuits following phase-dependent electrical stimulation of afferent pathways

**DOI:** 10.1101/2022.08.11.503603

**Authors:** Alejandro Pascual-Valdunciel, Nish Mohith Kurukuti, Cristina Montero-Pardo, Filipe Oliveira Barroso, José Luis Pons

## Abstract

Peripheral electrical stimulation (PES) of afferent pathways is a tool commonly used to induce neural adaptations in some neural disorders such as pathological tremor or stroke. However, the neuromodulatory effects of stimulation interventions synchronized with physiological activity (closed-loop strategies) have been scarcely researched in the upper-limb. Here, the short-term spinal effects of a 20-minute stimulation of afferent pathways protocol applied with a closed-loop strategy named Selective and Adaptive Timely Stimulation (SATS) was explored. The SATS strategy was applied to the radial nerve in-phase (INP) or out-of-phase (OOP) with respect to the muscle activity of the extensor carpi radialis (ECR). The neural adaptations at the spinal cord level were assessed for the flexor carpi radialis (FCR) by measuring disynaptic Group I inhibition, Ia presynaptic inhibition, and Ib facilitation from the H-reflex, and estimation of the neural drive before, immediately after, and 30 minutes after the intervention. SATS strategy was proved to deliver synchronous stimulation with the real-time measured muscle activity with an average delay of 17±8 ms. SATS-INP induced an increase of the disynaptic Group I inhibition (77±23 % of baseline conditioned FCR H-reflex), while SATS-OOP elicited the opposite effect (125±46 %). Not all the subjects maintained the changes after 30 minutes. Additionally, no other significant specific neural adaptations were found for the rest of measurements. These results suggest that the short-term modulatory effects of phase-dependent PES occur at the specific targeted spinal pathways for the wrist muscles in healthy individuals. Overall, timely recruitment of afferent pathways with the muscle activity is a fundamental principle which should be considered in tailoring PES protocols for the specific neural circuits to be modulated.

## 1. Introduction

Motor control is regulated by the central nervous system (CNS) through the interplay between afferent inputs from different sensory systems and supraspinal commands, which produces a coordinated and adaptive neural drive directed to the neuromusculoskeletal system [1]. By means of neural plasticity, synaptic transmission between neurons is potentiated or depressed in order to acquire or maintain stability of functions, a fundamental mechanism of learning and adaptation following neural injury or acquisition of new behaviors [2]. Plasticity at the spinal cord level plays a fundamental role in coordinating adaptive motor response to specific tasks [3]: interneurons are actively involved in the neuromodulation of afferent information, projecting it to the brain or directly to the spinal motoneurons and, therefore, adjusting the excitatory-inhibitory balance of specific pools of motor units [4].

Muscle spindles, which are stretch receptors, convey information on muscle length changes to the CNS through Ia afferent fibers, one of the main pathways involved in proprioception [5]. Particularly, the reciprocal inhibition pathway mediated by Ia fibers enables relaxation of the antagonist muscle to favor a functional movement. Nevertheless, some studies suggest that true reciprocal Ia inhibitory interneurons are replaced by the disynaptic Group I inhibitory interneurons, which receive major contribution from Ib fibers, and Ia fibers as well, and this mechanism mediates the inhibition of the antagonist muscle rather than the pure Ia pathway reported for the elbow or ankle joints [6].

There is still no consensus on whether short-term (from minutes to hours) spinal cord plasticity involves the CNS across different behaviors [7], or instead it is exclusively activity-dependent (i.e., plastic changes occur just at the neural circuitries involved in a specific motor task. Several studies have supported the idea that plastic changes are induced during interventions involving training of specific motor tasks [8]. Conversely, other studies have provided results supporting alterations of some CNS circuits even without engaging in any particular behavior or motor tasks [9,10].

In the past few years, peripheral electrical stimulation (PES) of afferent (sensory) pathways has been tested as a strategy to modulate the CNS aiming at restoring impaired functions in patients suffering from neural disorders such as spinal cord injury (SCI), stroke or essential tremor (ET) [11,12]. Specifically, some studies have applied PES to enhance activity-dependent plasticity of the CNS by recruiting afferent fibers paired with physiological activity [13]. For instance, patterned stimulation of tibialis anterior was shown to enhance reciprocal inhibition of the soleus in healthy subjects [14]. In another study, PES of the first dorsal interosseous muscle paired with single pulses of transcranial magnetic stimulation (TMS) at the corticospinal-motoneuron synapses was shown to improve corticospinal transmission in healthy and SCI patients [15]. Plastic changes induced after PES interventions are not only limited to neurophysiological measurements: functional improvements have been reported in different motor disorders [13]. Regarding pathological tremor, PES of afferent pathways targeting the antagonist muscle to the tremorgenic activity at the wrist level has been proved reduce pathological tremor in ET patients [16,17].

Despite the recent translation from bench to bedside of several PES-based therapeutic interventions in the upper-limb, including tremor management solutions, none of the prior studies has explored the short-term neural changes elicited after a PES intervention onto the muscles controlling the wrist [18]. The main goal of this study was to assess short-term spinal effects of a novel phase-dependent closed-loop PES strategy named Selective and Adaptive Timely Stimulation (SATS). To achieve this goal, the afferent fibers innervating extensor carpi radialis (ECR) were recruited by stimulating the radial nerve in-phase or out-of-phase with the muscle activity measured in real-time through EMG. This was done while healthy volunteers performed fast wrist flexion-extension movements (mimicking pathological tremor). A set of neurophysiological techniques including disynaptic Group I inhibition of the H-reflex, Ia presynaptic inhibition, Ib fibers facilitation and estimation of the neural drive, were applied to assess the effects of the intervention immediately after and 30 minutes after the stimulation ceased. It was hypothesized that in-phase and out-of-phase stimulation would elicit an increase and decrease in wrist disynaptic Group I inhibition of the H-reflex, respectively, similarly to what has been observed for the soleus and tibialis anterior muscles in the lower limb. By unravelling some of the spinal mechanisms underlying neural plasticity after time-dependent PES of afferent pathways, this study provides fundamental knowledge that can be used to develop therapeutic approaches for some neural impairments, including pathological tremor.

## 2. Methods

### 2.1. Participants

Fifteen healthy subjects volunteered to participate in the study. Eleven subjects (aged 22-27 years, 6 male, 5 female) completed the whole procedures, and four of them were excluded from the study since it was not possible to obtain a reproducible H-reflex from the FCR during the first experimental session. All procedures were approved by Shirley Ryan AbilityLab institutional review board (IRB protocol number STU00211930) and registered in Clinicaltrials.gov (NCT number 04501133). Participants signed a written informed consent in accordance with the Declaration of Helsinki.

### 2.2. Experimental paradigm

The short-term effects on FCR after the selective and adaptive timely stimulation (SATS) strategy applied in-phase (INP) or out-of-phase (OOP) with respect to the extensor carpi radialis (ECR) muscle activity were assessed (Figure 1-A). The study consisted of two experimental sessions for each participant, separated by at least 3 days (wash-out period). In each of the two sessions, one stimulation strategy randomly assigned (SATS-INP or SATS-OOP) was applied during the intervention phase. The subjects were blinded to the stimulation condition. The neurophysiological assessments were similar across visits and included three time points: baseline (PRE), immediately after the stimulation (POST), and 30 minutes after the stimulation (POST30’)

**Figure 1.**
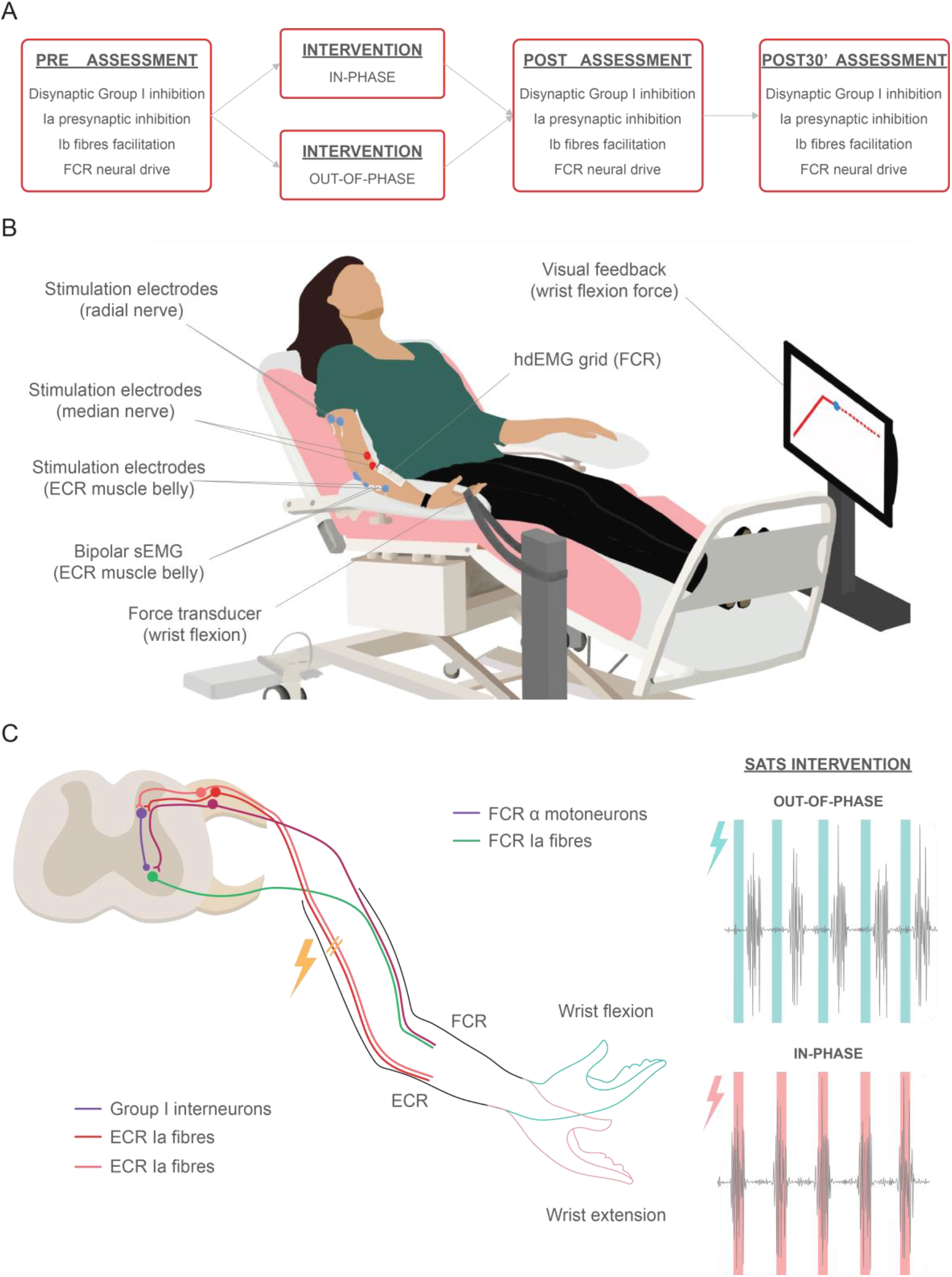
Experimental paradigm. A. Experimental protocol. Subjects underwent two experimental sessions in different days, in which the either SATS-INP or SATS-OOP strategy was applied for 20 minutes. Neurophysiological assessments were performed before, immediately after and 30 minutes after the intervention. B. Illustration of the experimental setup used in this study. C. Simplified schematics of the spinal circuits involved in the experimental hypothesis. On the right, grey lines represent EMG activity of the ECR muscle, while colored rectangles represent the applied stimulation. Note that during the SATS-INP intervention, the stimulation was delivered to the radial nerve when the ECR muscle was active, while during the SATS-OOP intervention, the stimulation was delivered out-of-phase to the ECR activity.

### 2.3. Experimental setup

Subjects comfortably sat on an armchair throughout the experimental procedures, during which they were asked to rest their dominant forearm on an armrest, keeping their shoulder extended and the elbow extended at approximately 45 degrees and 120 degrees in supination, respectively (Figure 1-B).

To record muscle activity during the neurophysiological assessments, high-density EMG (hdEMG) grids (13×5 electrode grid, 8 mm inter-electrode distance; OT Bioelettronica, Italy) were placed over the muscle belly of the flexor carpi radialis (FCR). Monopolar EMG signals were acquired at 2,048 Hz with a biosignal amplifier (Quattrocento; OT Bioelettronica, Italy) and stored for offline analysis. In addition to hdEMG assessments, bipolar surface (sEMG) electrodes were placed on the ECR muscle belly and the wrist flexors group (20 mm inter-electrode distance) in order to record muscle activity as input for the EMG-based stimulation strategies. A customized embedded system (OT Bioelettronica, Italy) with processing, stimulation and biosignal amplification capabilities was used to acquire the sEMG signals at 2,000 Hz. A wet wristband was used as reference for both EMG recordings modalities.

Two surface round stimulation electrodes (2 cm diameter, Axelgaard, Denmark) were placed over the median nerve at the cubital fossa, and two over the radial nerve at the spiral groove [6]. Prior to positioning the surface electrodes for stimulation, a stimulation bar was used to find the optimal location to elicit wrist extension when applying stimulation to the radial nerve, as well as to elicit wrist flexion and a reproducible FCR H-reflex when applying stimulation to the median nerve. Additionally, two bipolar surface stimulation electrodes (3.2 cm diameter, Axelgaard, Denmark) were placed over the ECR muscle belly (cathode proximal) to directly recruit the muscle fibers during the Ib fibers facilitation assessment. Two constant current stimulators (DS5, Digitimer, UK) were used to deliver the electrical stimuli during the neurophysiological assessments. The customized embedded system was used to run the stimulation strategies during the intervention.

During the neurophysiological assessments, a force sensor was used to measure the force exerted by the subject, which was displayed through visual feedback to guide the subject while performing the submaximal isometric flexion task (Figure 1-B).

### 2.4. Stimulation interventions

Subjects were asked to mimic typical tremorgenic movements by performing fast wrist flexion-extension with their dominant arm for 20 minutes. In each session, one of the two stimulation strategies (SATS-INP or SATS-OOP) was applied while subjects mimicked tremor movements (Figure 1-C). The intervention was divided into two periods of 10-minute stimulation each and 2 minutes of rest between them to minimize fatigue of the wrist muscles.

SATS consists of a closed-loop EMG-based stimulation in which tremorgenic bursts are detected in a pair of antagonist muscles through the EMG signals and stimulation is consequently applied to one of the muscles (antagonist or agonist) to activate afferent pathways timed with physiological activity (Figure 1-C). Stimulation was delivered through the surface electrodes placed on the radial nerve and a common ground electrode placed on the olecranon. Stimulation frequency was set to 100 Hz to optimize recruitment of Ia afferent fibers [19], pulse width was set to 400 μs [16], and duty cycle was set to 20 % of the tremor cycle to allow fast adaptation to rapid tremor movements. Stimulation intensity was calibrated for each subject prior to the intervention, being set immediately below motor and discomfort thresholds. SATS strategy operation was the following: 1 s recording windows were sequentially acquired and demodulated to determine the movement frequency. If the estimated frequency was within the tremor bandwidth (3-12 Hz), a 4 s stimulation window would be enabled. Along the stimulation window, the root mean square (RMS) of short EMG windows ∼15 ms) for each muscle were computed in real-time and compared to an adaptive threshold computed as the RMS during the recording window and multiplied by a gain factor. For the SATS-INP strategy, the radial nerve was stimulated when the simulated tremorgenic activity was detected in the ECR muscle (Figure 1-c). For the SATS-OOP strategy, the radial nerve was stimulated when the simulated tremorgenic activity was detected in the FCR muscle (Figure 1-C). After delivering a train of stimuli with a duration of 20 % of the tremor cycle, the RMS and stimulation epochs were sequentially repeated until the stimulation window was over. Then, the 1 s recording window was repeated again to assess the presence of tremor and to update the adaptive threshold to the tremor amplitude.

### 2.5. Neurophysiological assessments

#### 2.5.1. Disynaptic Group I inhibition and Ia presynaptic inhibition

Disynaptic Group I inhibition was assessed through the FCR H-reflex conditioning paradigm. Basal FCR H-reflexes were obtained by applying monopolar rectangular pulses of 1 ms over the median nerve [6]. The stimulation intensity of the test stimuli was set to elicit a FCR H-reflex with an amplitude on the ascending slope of the recruitment curve. The conditioned FCR H-reflexes were obtained by stimulating the radial nerve with 1 ms monopolar rectangular pulses prior (-1 ms), during (0 ms) or after (+1 ms) the application of the test stimuli [20]. The stimulation intensity of the conditioning stimuli was set immediately below motor threshold. For each condition, ten H-reflexes were elicited, with an inter stimulus interval (ISI) of 5 seconds to avoid post-activation depression [21]. The order of the test and conditioned H-reflexes was randomized. To facilitate the reflex response, subjects were asked to keep a slight wrist flexion during the assessment [20]. Additionally, presynaptic inhibition was measured following the same procedure by applying a conditioning stimulus on the radial nerve 20 ms prior to the test stimuli [22].

#### 2.5.2. Ib facilitation

A similar protocol to the one described by [23] was followed to assess the evoked ECR Ib fibers excitability. Stimulation of the ECR eliciting a visible tendon pull increases excitability of the FCR H-reflex by means of activation of Ib fibers and Ib interneurons. The FCR H-reflex was conditioned by applying a single stimulus (30 ms prior to the stimulation of the median nerve, 1 ms monopolar rectangular pulse) of the ECR muscle belly with an intensity 3 times above motor threshold. Ten unconditioned and ten conditioned FCR H-reflex responses were averaged to obtain representative measurements.

#### 2.5.3. Neural drive estimation

The excitability of the FCR motoneuron pool was assessed via hdEMG while the subjects performed a voluntary submaximal isometric contraction of the FCR muscle. At the start of the experiment, the subjects were asked to perform three maximum isometric voluntary contractions (MVC) of the FCR during 3 seconds while the elbow joint movement was constrained. The assessment consisted of one isometric wrist flexion at 10 % of the average maximal exerted force during the MCV trials. The contraction involved a 5-second ramp up, a 30-second plateau (10 % force of MVC) and a 5-second ramp down.

### 2.6. Data analysis

#### 2.6.1. SATS performance

The performance of the SATS strategy was estimated in order to demonstrate the phase difference between SATS-INP and SATS-OOP stimulation interventions. ECR and FCR sEMG signals were recorded during the intervention, and processed offline, in order to extract the components relative to the stimulation artefacts and mimicked tremor muscle activity. The isolation of the stimulation artefact component from the muscle activity was performed by applying a second order zero-phase Butterworth high-pass filter (500 Hz) to the raw sEMG signals, which resulted in the extraction of the stimulation and its harmonics. On the other hand, the muscle activity component was extracted from the raw signal by applying a Notch filter at 60 Hz to remove the power line interference, and a set of Notch filters to remove the contribution of the stimulation frequency and its harmonics (100 Hz, 200 Hz, 300 Hz, …, 900 Hz). Then, a second order zero-phase Butterworth band-pass filter (10-90 Hz) was applied to extract an estimation of the muscle activity component. Consequently, both muscle activity and stimulation components were band-pass filtered in the tremor band ±1 Hz with respect to the estimated mimicked tremor frequency in the power spectral density (PSD) computed for the ECR muscle [24]. The instantaneous phase was estimated by means of the imaginary part of the Hilbert transform [25]. The absolute phase difference between the muscle activity and the stimulation artefacts was computed and normalized to the range [0 2π]. Circular polar histograms with 20º bin resolution were produced with the phase difference between the muscle activity for ECR and FCR, and the SATS-INP and SATS-OOP stimulation artefacts [26]. Then, the absolute mean delay between the stimulation and the tremor activity was estimated based on the circular mean and the instantaneous frequency following a similar approach to that described in [27]. The ECR activity was selected as the reference muscle for computing phase difference and mean delays characterizing the stimulation performance. Ultimately, the phase difference and mean delay between the ECR and FCR mimicking tremor activity was estimated following the similar procedure for the OOP-SATS intervention in order to characterize the out-of-phase pattern.

#### 2.6.2. Disynaptic Group I inhibition, Ia presynaptic inhibition and Ib facilitation

FCR H-reflexes recorded via monopolar hdEMG were digitally filtered with a third order Butterworth band-pass filter (20-500 Hz). For each stimulation condition, ten responses to stimulation were averaged channel by channel. The intensity of the M-wave and H-reflex responses were estimated by computing peak to peak amplitude of the evoked potentials [14]. The hdEMG grid not only covered the FCR muscle belly, but also the surrounding muscles, thus, it was necessary to identify a region of interest (ROI) with the more prominent reflex activity elicited through the stimulation. Besides, though the same positioning protocol was followed for all the participants, the anatomic distribution of the muscles covered by the grid was variable across subjects [28]. Therefore, an automatic segmentation algorithm was applied to compute a two-dimensional activation map and determine a ROI for each condition. The matrix containing the average H-reflex amplitudes for each channel was used as input as a grayscale image to the algorithm based on the morphological segmentation techniques [29]. The resulting map was an umbralized or binary image containing the ROI of the H-reflex (Figure 2). The ROI defined for the basal or unconditioned H-reflex at the PRE assessment was used to extract the H-reflex response for the different conditioned and unconditioned H-reflex across time assessments (POST and POST30’). Ultimately, the channels within the ROI were averaged to provide the final H-reflex response. The conditioning time stimulus (-1 ms, 0 ms or 1ms) showing greatest inhibition of the FCR H-reflex at the PRE assessment was selected to assess the change in disynaptic Group I inhibition for the POST and POST30’ measurements.

**Figure 2.**
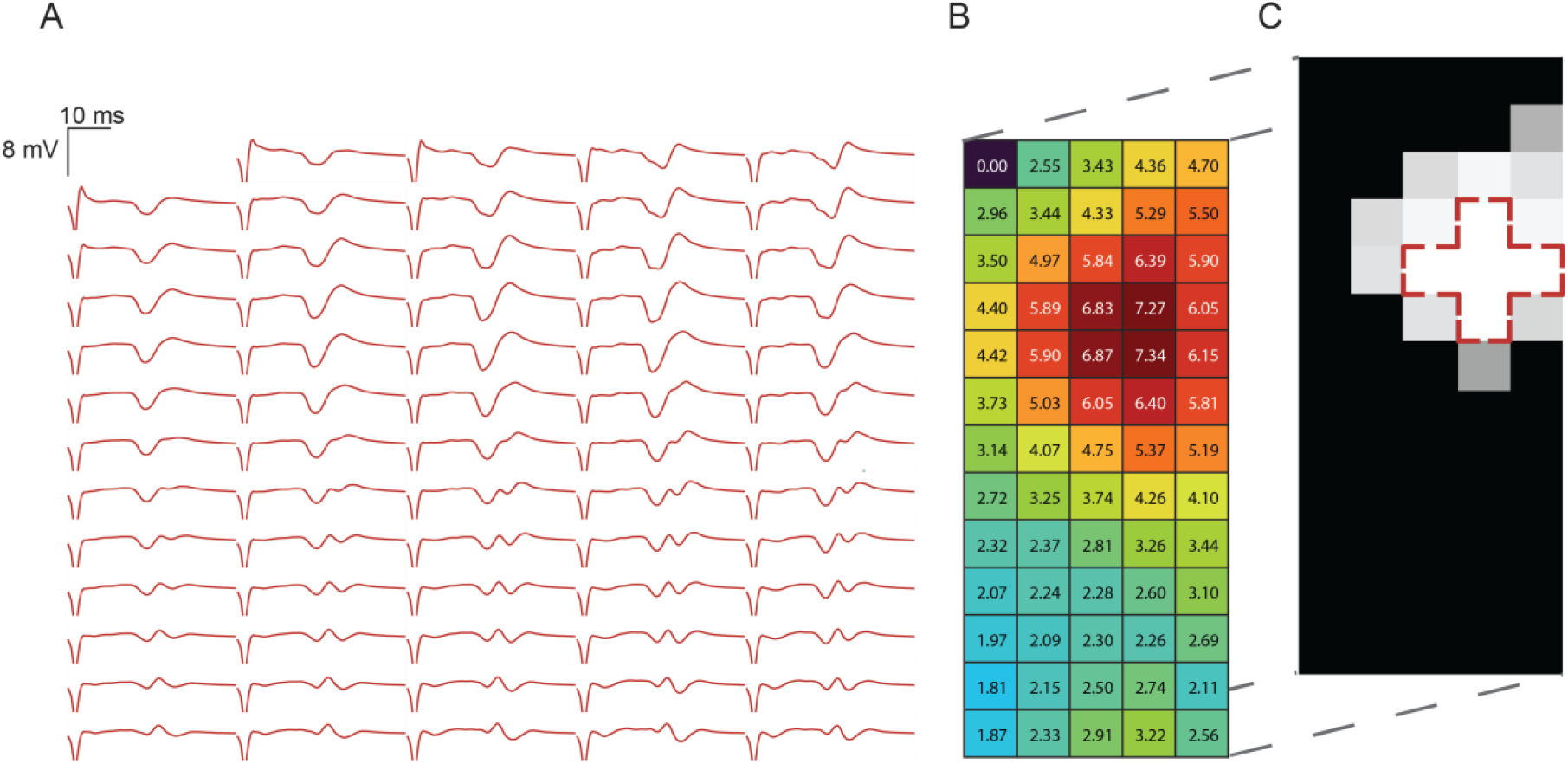
Illustration of the ROI segmentation of the basal H-reflex responses recorded for one of the subjects. A,B. The peak to peak amplitude of the FCR H-wave was estimated for each of the 64 channels. C. The resulting values were transformed into a grayscale image and a morphological segmentation method was applied to extract the ROI (red dashed lines).

The same analysis procedure described in the disynaptic Group I inhibition assessment was applied with the reflex responses recorded during the Ia presynaptic inhibition and Ib fibers facilitation assessment protocols.

#### 2.6.3. Neural drive estimation

Monopolar signals were digitally filtered with a second order Butterworth band-pass filter (20-500 Hz) and a Notch filter (60 Hz) to remove the power line interference. A convolutive blind-source separation algorithm was applied to decompose the monopolar signals into motor unit spike trains following the procedure described in [30]. Then, a manual inspection on the automatically identified spike trains was applied to remove duplicates, false positives and false negatives [31]. After manual edition, all the motor units (MUs) showing pulse-to-noise ratio above 30dB, which ensures sensitivity higher than 90 %, were kept for the subsequent analysis.

Instantaneous discharge rate was computed for each MU during the 30 seconds plateau of the submaximal contraction at 10 % force of the MVC by isolating the MU firing times associated with the plateau. Isolated MU firing times were used to compute inter-spike intervals, which were then smoothed and inverted to obtain instantaneous discharge rate, following the procedures described in [32]. Mean discharge rate for each MU was computed by averaging the instantaneous discharge rate for the given plateau.

The global EMG amplitude was used as an estimation of the neural drive [33]. The EMG filtered signals from the 64 hdEMG channels recorded during the MVC trials were segmented and the average RMS was computed by using a 250 ms moving window around 3 seconds containing the maximum peak of exerted force. The average RMS values for all channels were used as input to a segmentation algorithm to extract the ROI of the FCR, similar to the one used to extract the ROI for the FCR H-reflex [29]. Then, the EMG recorded during the 10 % MVC isometric wrist flexion task was normalized channel by channel to the RMS computed for the MVC task. The RMS values were computed over a 5 second window around the middle of the 30 s plateau, and the averaged RMS representing the EMG amplitude were computed for the channels contained within the ROI.

The force produced by the subjects during the submaximal wrist flexion was also analyzed. Similar to the analysis of the EMG amplitude, the force signals were normalized to the maximum force produced during the MVC task. A representation of the force level was estimated by means of the RMS applied over a 5 seconds window around the middle of the 30 seconds plateau. Alternatively, the PSD of the 30 s plateau was estimated using Welch’s periodogram method. The power of the force exerted corresponding to the frequency band related to the frequency band of the mimicked pathological tremor and physiological tremor was estimated by averaging the PSD in the [3-6] Hz and [5-12] Hz frequency bands, respectively [34]. The goal of this analysis was to explore the potential effects of the phase-dependent stimulation in the specific frequency bands of motor control related to the stimulation applied and the physiological tremor.

#### 2.6.4. Statistical analysis

All statistical analyses were performed using linear mixed effect models in R. Post-hoc comparisons between assessments (PRE, POST, POST30’) and stimulation phase (SATS-INP or SATS-OOP) were applied using Holm-Bonferroni corrected t-tests. Distributions were inspected for normality for all the data examined here. Firstly, it was evaluated whether there was a significant difference between conditions for different phases. Then a mixed model analysis was performed of the disynaptic Group I inhibition, Ia presynaptic inhibition, Ib facilitation and mean discharge rate of the MUs during isometric wrist flexion with fixed effects of condition, phase, and interaction between condition and phase, and random slopes and intercepts for each subject. Paired t-tests were also applied to investigate the frequency difference for the mimicked tremor movements and the phase difference between the stimulation and muscle activity between experimental sessions.

## 3. Results

A total of 11 subjects completed both experimental sessions (SATS-INP and SATS-OOP). All the stimulation interventions were delivered in the range between 4 and 9 mA (always below motor threshold), with no adverse effects being reported by any of the subjects. Subjects were able to perform rapid flexion-extension wrist movements mimicking tremor, with the estimated frequency for ECR muscle activity being similar during both phase-dependent interventions (p<0.05; 4.3±0.5 Hz for the SATS-INP intervention, and 4.4±0.7 Hz for the SATS-OOP intervention).

### 3.1. SATS performance

Figures 3-A,B show the circular histograms with the phase difference values (degrees) between the stimulation and the ECR muscle activity for SATS-INP and SATS-OOP interventions. The absolute mean phase difference between the ECR activity and the stimulation applied across subjects for SATS-INP intervention was 26±10 degrees (equivalent to a delay of stimulation of 17±8 ms), while for SATS-OOP, the absolute mean phase difference was 127±29 degrees (equivalent to a delay of 81±21 ms). These results showed that the applied stimulation strategy was different (p<0.05) in both interventions. The delay results between the stimulation timing and the ECR muscle activity for the SATS-OOP strategy were not completely aligned with an out-of-phase pattern of 180 degrees. Therefore, the average phase difference and delay between the ECR and FCR muscle activity was estimated in order to characterize the physiological out-of-phase pattern of mimicked tremor. Mean phase difference between ECR and FCR was 155±14 degrees, equivalent to 98±18 ms delay (Figure 3-C).

**Figure 3.**
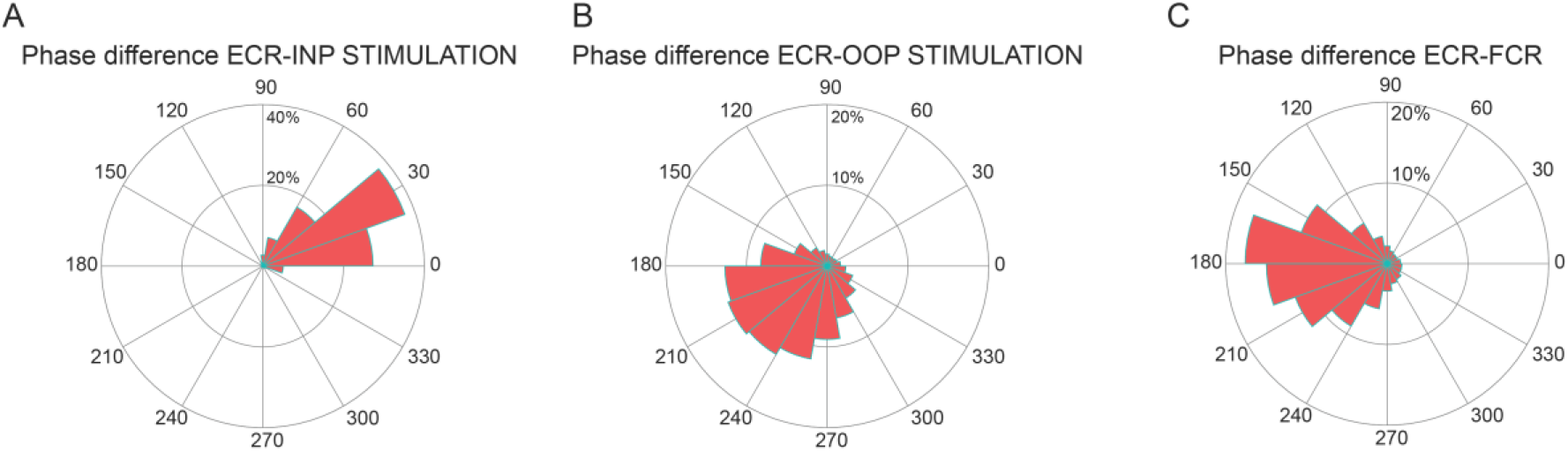
Circular histograms containing the estimated phase differences. The size of the bins represents the cumulative count of bins representing the phase difference (degrees) normalized between 0 and 100 % from all the subjects. A. Phase difference between the mimicked tremor activity of the ECR and the stimuli trains delivered using the SATS-INP strategy. B. Phase difference between the mimicked tremor activity of the ECR and the stimuli trains delivered using the SATS-OOP strategy. C. Phase difference between the mimicked tremor activity of the ECR and FCR during the SATS-OOP interventions.

### 3.2. Disynaptic Group I inhibition, Ia presynaptic inhibition and Ib facilitation

Figure 2 illustrates the procedure followed to extract the average FCR H-reflex peak to peak amplitude from the hdEMG grid for each set of conditioning stimuli. Basal (unconditioned) FCR H-reflex at the PRE assessment was on average 57±62 % and 50±64 % of maximal M-wave for the SATS-INP and SATS-OOP sessions respectively. Neither SATS-INP nor SATS-OOP elicited statistically significant changes in the basal FCR H-reflex amplitude at the POST and POST30’ assessment (fixed effects: stimulation phase p=0.92; time p=0.55; interaction, p=0.59), reflecting that the reflex response remained stable across time and interventions.

Figures 4-A,B,C,D illustrate the disynaptic Group I inhibition measured at the conditioned H-reflex for one of the subjects during the SATS-INP experimental session. Average disynaptic Group I inhibition increased at POST (77±23 % of conditioned FCR H-reflex at PRE) and POST30’ (84±18 % of conditioned FCR H-reflex at PRE after the SATS-INP intervention, while the SATS-OOP intervention elicited the opposite effect, resulting in decreased inhibition at POST (125±46 % of conditioned FCR H-reflex at PRE), but not at POST30’ (101±26 % of conditioned FCR H-reflex at PRE) (Figures 4-E,F). Linear mixed effect analysis for disynaptic Group I inhibition found that there was no significant main effect of the time assessment (p = 0.529) and stimulation phase (p = 0.051), but found a significant main effect of the interaction between time assessment and stimulation phase (p < 0.001). The post-hoc comparisons revealed statistically significant differences between SATS-INP POST and SATS-OOP POST (p < 0.05), and SATS-INP PRE and SATS-INP POST (p < 0.05), while there were no significant differences between any of the other comparisons.

**Figure 4.**
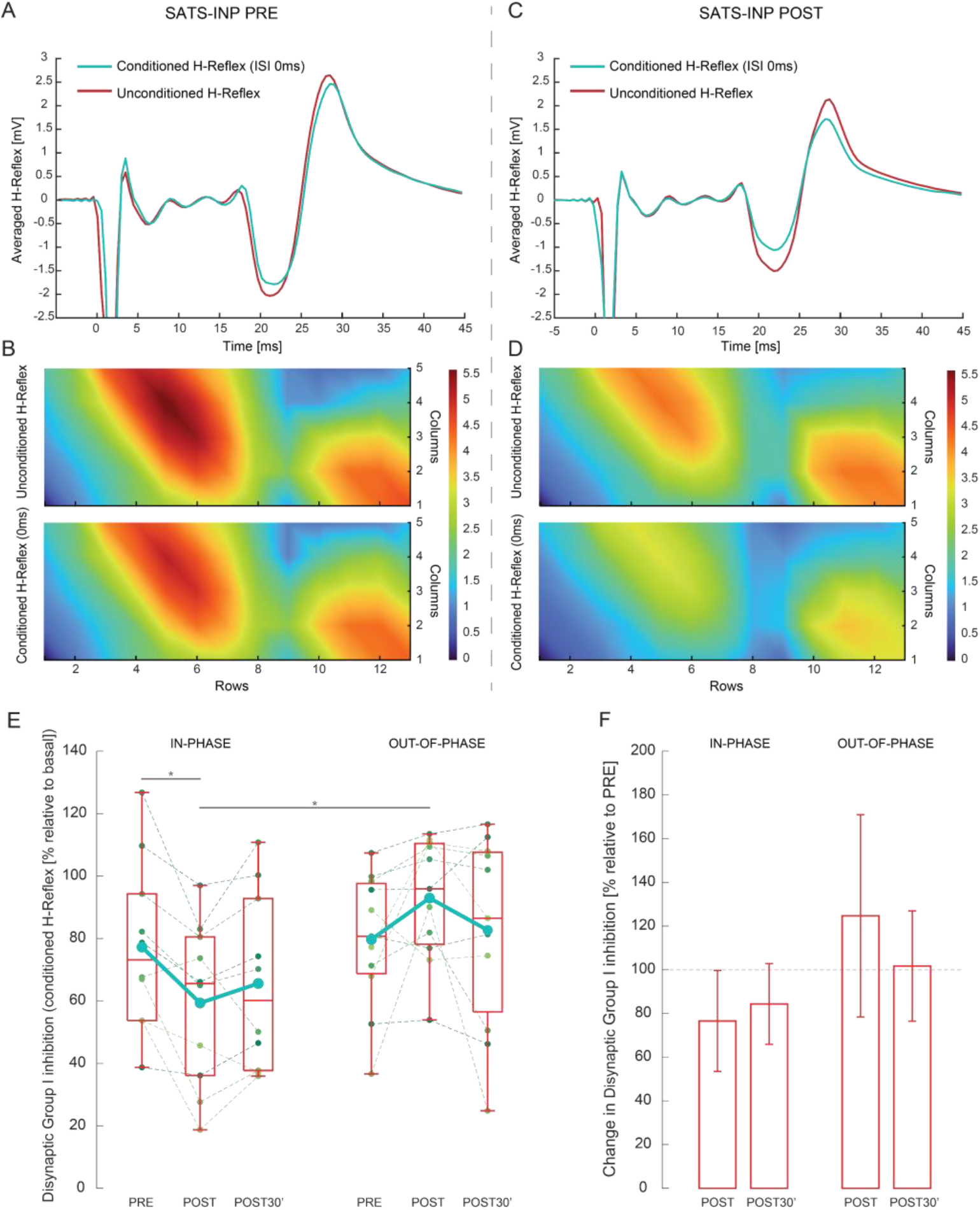
Disynaptic Group I inhibition measured with the FCR H-reflex conditioning paradigm. A-D. Example of FCR H-reflex responses of one of the subjects. A, C. average of ten baseline and ten conditioned FCR H-reflex responses of the ROI recorded at the PRE and POST assessments during the SATS-INP session, respectively. B, D. 2D colormap representing the peak-to-peak FCR H-reflex amplitude through the hdEMG grid. E. Boxplots representing the disynaptic Group I inhibition across time assessments and stimulation conditions. Small dots and dashed lines represent individual subject data. Solid green lines represent averaged values across subjects, while red horizontal lines within the boxes represent the median values. F. Average change and standard deviation in disynaptic Group I inhibition at the POST and POST30’ assessments compared to the PRE assessment. * p-value < 0.05; Holm-Bonferroni corrected t-test.

Ia presynaptic inhibition measured through the conditioning of FCR H-reflex with a stimulus delivered at the radial nerve 20 ms prior to the test stimulus was not modified by any of the two interventions (Figures 5-A,B). Following SATS-INP intervention, average Ia presynaptic inhibition resulted in 99±27 % and 94±28 % of the PRE values for the POST and POST30’ assessments, respectively, while following SATS-OOP intervention, it resulted in 107±29 % and 117±39 % of the PRE values for the POST and POST30’ assessments, respectively. Linear mixed effect analysis found no significant main effect of the time assessment (p = 0.814), stimulation phase (p = 0.455), and the interaction between both factors (p = 0.291).

**Figure 5.**
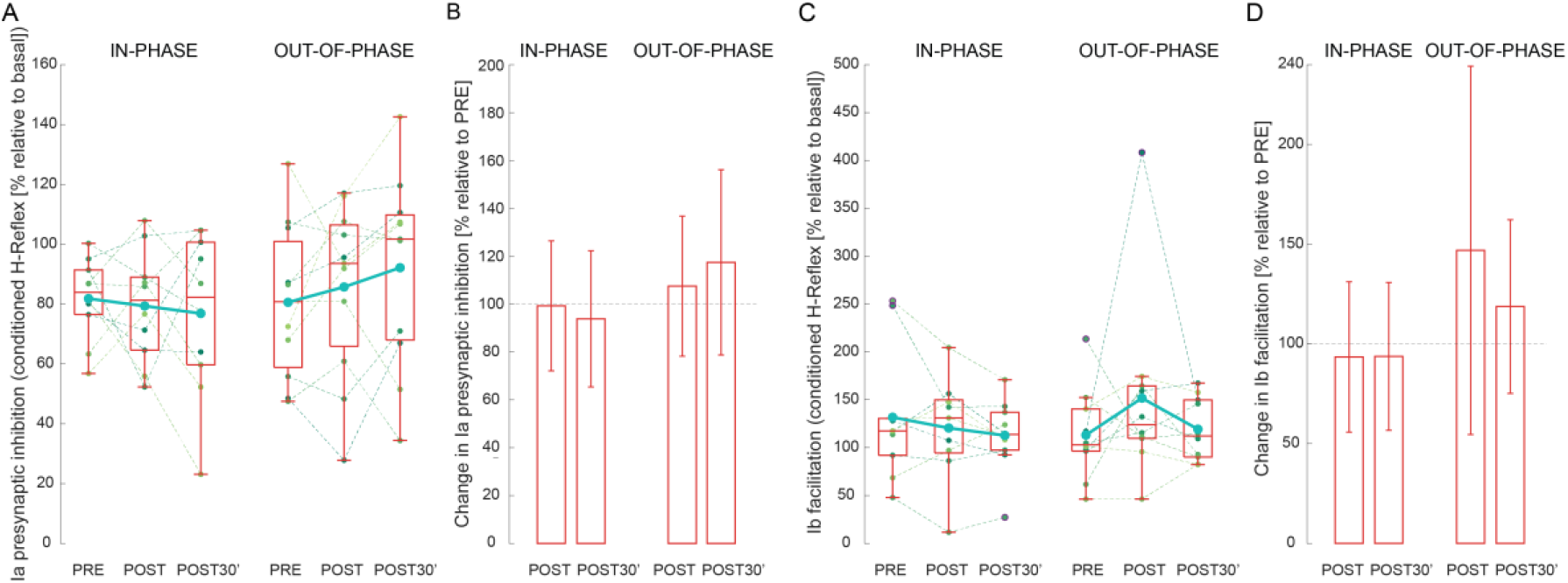
Ia presynaptic inhibition and Ib facilitation. A, C. boxplots representing Ia presynaptic inhibition and Ib facilitation across time assessments and stimulation conditions, respectively. Small dots and dashed lines represent individual subject data; solid green lines represent averaged values across subjects; red horizontal lines within the boxes represent the median values. B, D. average change and standard deviation in Ia presynaptic inhibition and Ib facilitation at the POST and POST30’ assessments compared to the PRE assessment, respectively.

SATS-OOP intervention elicited an increase of average Ib facilitation for the POST (147±92 % of conditioned H-reflex at PRE) and POST30’ (111±57 % of conditioned H-reflex at PRE) assessments compared to the PRE assessment, while for the SATS-INP, the mean Ib facilitation was 93±38 % (POST) and 94±37 % (POST30’) compared to the PRE assessment (Figures 5-C,D). However, due to the great variability the linear mixed effect analysis for Ib facilitation found no significant main effect of the time assessment (p = 0.768), stimulation phase (p = 0.880), and the interaction between time assessment and phase (p = 0.414)

### 3.3. Neural drive estimation

Neither SATS-INP nor SATS-OOP interventions induced changes in FCR neural drive estimated through global EMG amplitude, as shown in Figures 6-A,B (fixed effects: stimulation phase p=0.863; time p=0.420; interaction, p=0.273). Similarly, the force level production remained unchanged during the isometric wrist flexion task after applying both stimulation interventions (fixed effects: stimulation phase p=0.241; time p=0.064; interaction, p=0.938), meaning the submaximal wrist flexion task protocol was reproducible across time assessments (Figures 6-C,D). The linear mixed models did not show any significant effect in the power of the force produced in the stimulation frequency band (fixed effects: stimulation phase p=0.898; time p=0.245; interaction, p=0.842, Figures 6-E,F) or in the power of the physiological tremor frequency band (fixed effects: stimulation phase p=0.938; time p=0.943; interaction, p=0.953, Figures 6-G,H). It is noteworthy that in 18 out of 22 experimental sessions, the spectral power of the force produced in the stimulation frequency band was increased after the intervention (Figure 6-E).

**Figure 6.**
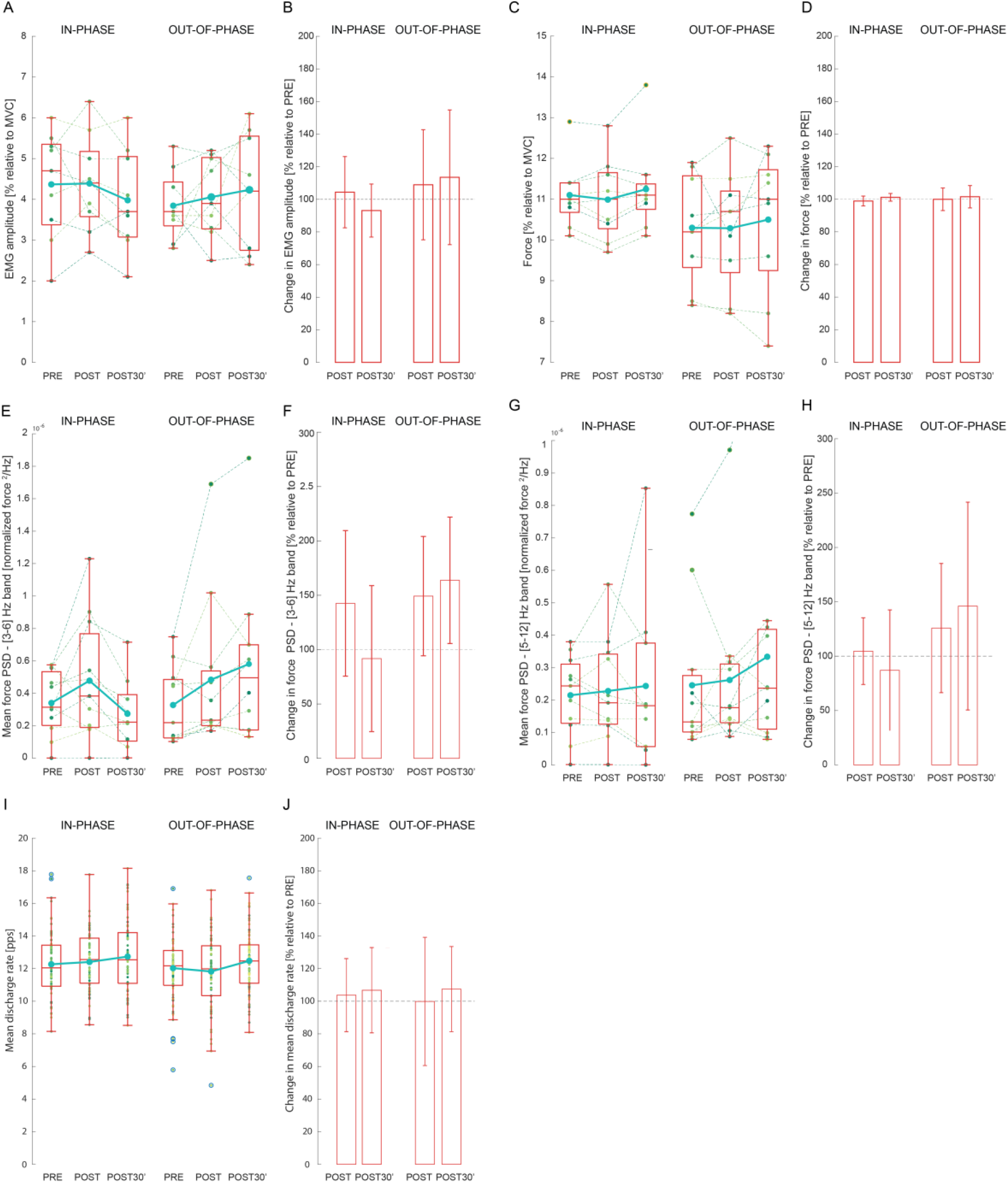
Neurophysiological assessments of the neural drive. A, C, E, G, I. Boxplots representing EMG amplitude, level of force produced, power of the force produced in the [3-6] Hz and [5-12] Hz bands, and MUs mean discharge rates across time assessments and stimulation conditions, respectively. A, C, E, G. Small dots and dashed lines represent individual subject data; solid green lines represent averaged values across subjects; red horizontal lines within the boxes represent the median values. I. Small dots represent single identified MUs; solid green lines represent averaged values across subjects; red horizontal lines within the boxes represent the median values. A, C, E, G, I. solid green lines represent averaged values across subjects. B, D, F, H, J. Average change and standard deviation in EMG amplitude, level of force produced, power of the force produced in the [3-6] Hz and [5-12] Hz bands, and MUs mean discharge rates at the POST and POST30’ assessments compared to the PRE assessment, respectively.

The neural drive estimated through the MUs mean discharge rate during the plateau of the isometric wrist flexion task did not change across time due to either SATS-INP or SATS-OOP interventions, as shown in Figures 6-I,J (fixed effects: stimulation phase p=0.569; time p=0.082; interaction p=0.296).

## 4. Discussion

The main goal of this study was to assess short-term spinal effects of applying phase-dependent PES. To achieve this goal, the radial nerve was stimulated in-phase or out-of-phase with the ECR muscle activity while healthy volunteers performed fast wrist flexion-extension movements (mimicking pathological tremor). Neurophysiological changes were assessed before, immediately after and 30 minutes after stimulation trials. Results supported the hypothesis that in-phase and out-of-phase stimulation would elicit an increase and decrease in disynaptic Group I inhibition, respectively. Further assessments measuring specific neural circuits such as Ib fibers facilitation, Ia presynaptic inhibition or the estimation of the neural drive during submaximal flexion task did not provide robust evidence of the presence of additional neural mechanisms altered after the interventions. These results suggest that the neural adaptations derived from the stimulation of the antagonist muscle might be specific to the predominant spinal circuit activated (disynaptic Group I inhibition) and modulated during the task performed.

The hand is the end effector of the upper limb and requires fine motor control, which is ultimately achieved by means of synergistic activation of the intrinsic and extrinsic extensors and flexors of the hand [35]. Thus, pure agonist-antagonist muscle behavior might be limited to a few motor tasks, being the rapid and cyclic wrist flexion-extension contractions among them [6]. It has been hypothesized that the fusimotor system adapts the gain of the muscle spindles to the stretching of the antagonist muscle [36]. Therefore, the PES intervention here tested was delivered while the subjects performed voluntary rapid flexion-extension movements as if they were mimicking hand tremor. The selection of this task allowed the activation of the wrist flexor and extensor groups as antagonist muscles, and the rapid movements with an average frequency around 4 Hz increased the amount of trains of electrical stimuli paired with the muscle activity, favoring the induction of plasticity mechanisms based on Hebbian principles [37]. The mean delay between the stimulation artefacts and the mimicked tremor muscle activity was also investigated to test the accuracy of the SATS strategy. Results showed that the subjects were capable of performing the alternating contractions at the required frequency and following a FCR-ECR activation nearly similar to the out-of-phase pattern described for patients exhibiting pathological tremor [27]. Likewise, the analysis showed that the SATS strategy locked the stimulation to the phase of the muscle activity with an average delay below 17 ms. Considering that the RMS windows used to detect real-time muscle activity were set to 15 ms, the SATS strategy provided accurate stimulation synchronously with the muscle activity.

### 4.1. Identification of the adapted neural mechanisms

Previous studies in the lower limb showed that the repetitive delivery of electrical stimuli to the common peroneal nerve induced an increase in the Ia reciprocal inhibition for the soleus muscle if the stimulation was synchronized with the swing phase of gait, while the opposite effect was induced when the stimulation was applied during the stance phase [38]. This study expands those findings to the upper-limb and the disynaptic Group I inhibition circuit at the wrist. Modulation of the wrist Group I inhibitory interneurons was dependent on the phase of stimulation. The tested protocol induced neuromodulation of the aforementioned circuits. However, they were not effectively maintained 30 minutes after the application, similarly to other studies applying stimulation protocols paired with physiological activity in the lower limb [14,39]. Nevertheless, it is noteworthy that in ten out of eleven subjects the disynaptic Group I inhibition values remained reduced compared to the baseline levels 30 minutes after the stimulation with the SATS-INP protocol. While changes in reciprocal inhibition mentioned in other studies have been attributed to modulation of Ia fibers or Ia interneurons for the ankle [38], the absence (or difficulty to demonstrate) true Ia reciprocal inhibition for the wrist muscles prevents us from assuming that similar changes occurred in one or more sites along the spinal circuit in this study [6]. Additional measurements of Ia presynaptic inhibition assessment did not reveal significant modulation of this mechanism, although descriptive statistics showed a similar trend to the disynaptic Group I inhibition, with an increase of Ia presynaptic inhibition after the stimulation was delivered in-phase with ECR activity. The presented evidence is not sufficient to determine the short-term modulation of Ia fibers, either at presynaptic or postsynaptic level into the disynaptic Group I interneurons. Regarding the excitability of FCR motoneurons and Ia fibers, no significant alterations of the basal FCR H-reflex were found after the interventions, which suggests that excitability of Ia fibers and motoneurons from FCR were affected by neither the phase of the stimulation nor the task performed.

Disynaptic Group I inhibition is mediated by multiple afferent fibers arising from homonymous, heteronymous and antagonist muscles forearm muscles, including Ib fibers, as well as Ia and possibly other group II afferents [6]. Hence, excitability of Ib fibers was non-invasively assessed to describe the contribution of these afferents to the disynaptic Group I inhibition modulation [23]. According to these results, the measurement of the excitability of Ib fibers was not altered after the stimulation interventions, although results should be carefully interpreted due to the high variability reported. The compilation of neurophysiological assessments supports the hypothesis that short-term modulation induced after SATS interventions occurs at the disynaptic group I inhibitory interneurons.

Measurements based on evoked potentials (e.g. H-Reflex or reciprocal inhibition tests) might be insufficient to describe the extent of the neural changes after an intervention. Certainly, assessment of the motoneuron properties can be achieved by estimation of the neural drive, which represents the synaptic input to the motoneurons decoded in spike trains sent to the muscles [33]. High density EMG (hdEMG) allows reliable and non-invasive estimation of the neural drive by means of surface EMG decomposition into motor unit spike trains [40]. Analysis of a pool of motor unit discharge rate properties provides valuable information about the neural control of the active pool of motoneurons at the spinal cord [41]. Consequently, complementary assessments of the neural drive while the subjects performed submaximal isometric flexion of the wrist sought to determine whether the neural adaptations were limited to the targeted spinal circuit or could be expanded to other behaviors of motor control. Neither the force level produced nor the neural drive estimated through normalized EMG amplitude were altered after the interventions. When analyzing the neural drive estimation by means of differences in MU discharge rate, results showed that MU discharge rate behavior remained consistent after applying the two interventions (SATS-OOF or SATS-INP), suggesting that none of the stimulation patterns induced significant effects on the MU discharge rate.

Particularly, power spectral analysis of the force produced during the submaximal wrist flexion revealed a potential increase of the power in the frequency band associated with the stimulation after both phase-locked interventions. These changes could be attributed to an oscillatory disruption in the stable motoneuron recruitment elicited by the stimulation delivered at a given frequency, which was proven to alter spinal circuits through feedback inputs from afferent fibers. Since the changes were reported after both interventions using opposite phase-locked stimulation, the execution of the rapid flexion-extension at the same frequency could also be responsible for the neural adaptations specific to the frequency band.

Overall, the timely recruitment of the afferent fibers has been proved to be determinant to induce specific neural adaptations at the spinal cord. Particularly, Ia or Ib fibers were not found to be conclusively modulated by the stimulation. Instead, disynaptic Group I interneurons are proposed as the site where short-term modulation may occur. However, these phase-specific neural adaptations located at the spinal cord cannot be attributed to be exclusively mediated by local spinal modulation mechanisms: other supraspinal pathways may instead be involved in the neuromodulation [42,43]. The changes in disynaptic Group I inhibition were assessed through the FCR H-reflex conditioning paradigm, and therefore those alterations could represent the global excitability of the CNS [44]. On the other hand, the outcomes from the neural drive assessments support the evidence that short-term neuromodulation could be specific to the wrist flexion-extension task performed and no major alterations of the CNS were elicited [7]. Then, it remains unresolved whether other mechanisms such as the propriospinal system along with other supraspinal circuits are adapted after phase-locked stimulation.

### 4.2. Limitations of the study

Despite the single-blinded paired randomized study design, the relatively reduced sample size (eleven subjects) limits the generalization of the results. Further studies with larger sample sizes can add more evidence and enlighten the source and duration of the short-term neural adaptations.

The average duration of each experimental session was approximately 3 hours, from which all the neurophysiological assessments were performed following the same order. The assessment of disynaptic Group I inhibition was performed in the first place to prioritize these outcomes. It cannot be ruled out that the different assessments, such as the Ib fibers excitability, might be affected by the fatigue induced as a result of the length of the experiment, as well as by the total amount of test and conditioning electrical stimuli applied over the radial and median nerves during the FCR H-reflex assessment trials.

The study was designed to characterize the neural adaptations of FCR motor control elicited after a PES-based intervention. Alterations of the ECR derived from the stimulation of the ECR were not tracked during the experiments. However, information about the excitability of the ECR motoneurons, as well as the Ia and Ib fibers arising from the ECR would support the identification of the neural mechanisms modulated based on the stimulation phase. Moreover, addition of measurements aiming at identifying the potential alterations on brain circuits and their projections into motoneurons and spinal interneurons will be necessary to fully clarify the extent of the neuromodulatory effects across the CNS.

### 4.3. Implications for neural rehabilitation and tremor reduction

This is the first study exploring the short-term neuromodulatory effects on the FCR after an intervention using activity phase-locked PES of afferent pathways. The outcomes of this study with healthy subjects will contribute not only to a better understanding of the physiology of the spinal circuits controlling the wrist, but also to describe some of the neural adaptations elicited after PES interventions used as a neural rehabilitation tool in different motor disorders such as pathological tremor or spasticity. In the case of ET, alternating activation of agonist and antagonist muscles of the wrist and the contribution of Ia fibers have been hypothesized to favor pathological tremor through a mechanism similar to Ia reciprocal inhibition [45]. Additional studies in favor of the implication of afferent fibers in pathological tremor suggest that cutaneous and Ib fibers are involved in both tremor and rigidity in Parkinson’s disease through the propriospinal system [46,47]. On the other hand, dysregulation of spinal pathways such as the stretch reflex, or the Ia reciprocal and presynaptic inhibitions, among others, might play a major role in spasticity [48,49].

In those scenarios where the spinal cord receives aberrant input from supraspinal pathological circuits, which are then amplified through afferent loops, the timing of PES could be used to specifically modulate spinal pathways by shifting the net excitatory output projected to the muscles towards functional states. Moreover, there is evidence of other forms of sensory PES targeting and modulating brain structures with functional implications [18]. Thus, the phase-locked strategies explored here could presumably induce other activity-dependent plastic mechanisms in multiple sites along the CNS.

## Acknowledgements

This study was funded by the European Union’s Horizon 2020 research and innovation program (EXTEND—Bidirectional Hyper-Connected Neural System, agreement N° 779982).

